# Chemical transfection reagents induce intracellular Ca^2+^ signals in TRPA1-expressing cells

**DOI:** 10.64898/2026.09.16.751488

**Authors:** Alina Milici, Andrei Segal, Justyna B. Startek, Karel Talavera

## Abstract

Transient receptor potential ankyrin 1 (TRPA1) is a polymodal sensory ion channel whose activity is influenced not only by chemical ligands but also by the physical properties of the plasma membrane. This raises the possibility that membrane-active compounds used routinely in cell biology may alter TRPA1 function. We investigated the acute effects of two widely used chemical transfection reagents, Lipofectamine 3000 and Mirus *Trans*IT-293, on intracellular Ca²⁺ signaling and TRPA1 activity. For this, we monitored intracellular Ca²⁺ dynamics using ratiometric Fura-2 imaging in CHO cells stably expressing mouse TRPA1 (CHO-mTRPA1), parental CHO-WT cells, and primary mouse dorsal root ganglion (DRG) neurons. Transfection reagent preparations were applied at different concentrations under controlled temperature and low-flow conditions. The contribution of TRPA1 and Ca²⁺ influx was assessed using the selective TRPA1 inhibitor HC-030031 and the broad-spectrum Ca²⁺ channel blocker ruthenium red. We found that Lipofectamine 3000 induced concentration-dependent, irregular Ca²⁺ transients in CHO-mTRPA1 cells, while simultaneously inhibiting the constitutive TRPA1-dependent Ca²⁺ activity observed under basal conditions. Its inhibitory effect was evident at concentrations below those producing substantial cellular activation and was rapidly reversible after washout. At higher concentrations, Lipofectamine-induced Ca²⁺ responses were only partially suppressed by TRPA1 inhibition, indicating the involvement of additional mechanisms. Consistent with this, Lipofectamine also induced Ca²⁺ transients in CHO-WT cells and primary DRG neurons, where both extracellular Ca²⁺ influx and intracellular Ca²⁺ mobilization contributed to the responses. Analysis of the components of the Lipofectamine 3000 formulation further revealed distinct effects of Lipofectamine and the P3000 enhancer. Mirus *Trans*IT-293 similarly induced Ca²⁺ transients in CHO-mTRPA1 cells and DRG neurons. In CHO-mTRPA1 cells, its response was concentration-dependent and strongly reduced by HC-030031, whereas the response in DRG neurons showed little sensitivity to TRPA1 inhibition. We conclude that chemical transfection reagents can acutely alter intracellular Ca²⁺ homeostasis and modulate TRPA1 activity. Their effects involve both TRPA1-dependent and TRPA1-independent mechanisms and differ substantially between formulations and cell types. These findings identify membrane-active transfection reagents as previously underappreciated modulators of sensory ion-channel function and highlight their potential to influence the interpretation of experiments performed in transfected cells.

## Introduction

The ability of cells and organisms to respond to changes in their environment relies on molecular systems capable of detecting a broad range of external stimuli. Among these systems, Transient Receptor Potential (TRP) channels constitute a diverse family of plasma membrane proteins that respond to chemical, thermal and mechanical cues [1–5]. TRPA1, the only mammalian member of the ankyrin subfamily, is one of the most broadly tuned TRP channels and is prominently expressed in primary sensory neurons, where it contributes to the detection of noxious environmental stimuli and to pain signaling [6,7].

Although TRP channels can be activated by specific ligands, increasing evidence indicates that their activity is also strongly influenced by the physical properties of the surrounding plasma membrane. In particular, TRPA1 has been implicated as a mechanosensitive channel and can be modulated by changes in membrane lipid composition and organization compounds [4,8–11]. We previously showed that several chemical compounds can activate TRPA1 through mechanisms associated with mechanical perturbations of the plasma membrane rather than through classical ligand–channel interactions alkylphenols [10]. Similarly, bacterial lipopolysaccharides (LPS) have been proposed to activate TRPA1 by inserting into the lipid bilayer, thereby altering membrane organization and producing local mechanical effects on the channel [11]. The activity of TRPA1 is also sensitive to cholesterol-dependent membrane organization, including changes affecting lipid raft-associated structures by agonists [12–14]. Together, these observations support the view that TRPA1 can function not only as a receptor for chemically reactive molecules but also as a broadly tuned sensor of changes in the physical state of its membrane lipid environment [15].

This property raises an important question concerning the effects of exogenous lipid-based materials routinely introduced into experimental cells. Transfection reagents are widely used in cell biology to facilitate the intracellular delivery of nucleic acids. Among the most commonly employed reagents are lipid-based formulations such as Lipofectamine (lipoplex-forming) and Mirus (polyplex-forming), which promote the association of nucleic acids with lipid assemblies and facilitate their interaction with the plasma membrane, thereby enabling cellular uptake and subsequent delivery of their cargo. Their biological function therefore intrinsically depends on interactions with cellular membranes, involving processes such as adsorption to the plasma membrane, membrane reorganization, endocytosis and, depending on the formulation and experimental conditions, membrane fusion or disruption.

Transfection reagents are generally considered experimental tools whose effects are interpreted primarily in terms of their ability to improve transfection efficiency. However, because they are deliberately designed to interact with cellular membranes, their presence may also modify membrane properties and the activity of membrane proteins independently of the intended nucleic-acid delivery process. Such effects are particularly relevant for sensory ion channels whose gating is coupled to the physical state of the plasma membrane. Acute modulation of membrane proteins by lipid-based delivery systems could consequently represent an underappreciated source of variability in experiments involving transfected cells.

TRPA1 provides a particularly suitable model in which to investigate this possibility. Its sensitivity to membrane composition and mechanical perturbations suggests that compounds or supramolecular assemblies capable of interacting with the plasma membrane may influence channel activity even in the absence of a specific chemical interaction with the protein. This consideration is especially important in heterologous expression experiments, in which TRPA1 is commonly introduced into cells using lipid-based transfection reagents. If these reagents themselves alter TRPA1 function, they could influence the interpretation of experiments in which channel activity is subsequently assessed, potentially affecting measurements of agonist responses, channel sensitivity or downstream cellular signaling.

Despite the widespread use of lipid-based transfection reagents, their acute effects on ion-channel function have received comparatively little attention. In particular, whether commonly used transfection formulations can directly modulate the activity of TRPA1 has not, to our knowledge, been systematically investigated. Here, we addressed this question by examining the effects of two widely used lipid-based transfection reagents, Lipofectamine and Mirus, on TRPA1 function. By assessing channel activity following exposure to these reagents, we investigated whether membrane-interacting transfection formulations can acutely alter TRPA1 function and thereby reveal an additional dimension of the interaction between membrane-active compounds and sensory ion channels.

## Materials and methods

### Cell culture

Chinese hamster ovary cells (RRID:CVCL_0213) (denoted CHO-WT) from the American Type Culture Collection were grown in DMEM containing 10% fetal bovine serum, 2% glutamax (Gibco/Invitrogen), 1% non-essential amino acids (Invitrogen) and 200 µg/mL penicillin/streptomycin at 37 °C in a humidity-controlled incubator with 5% CO_2_. As mTRPA1 expression system we used CHO-K1 cells (RRID:CVCL_0214) stably transfected with mTRPA1 (denoted CHO-mTRPA1). The cells have been seeded on poly-L-lysine-coated 13 mm coverslips 2h before the measurements and cultured in 24-well plates.

### Microinjection system and temperature control

We implemented several modifications in the setup to replicate more closely the *in vivo* scenario of slow particle diffusion through the tissue. We replaced the classic, constant perfusion of the cells with a gentle injection of test solutions,

For this, we used a microinjection syringe pump (Fusion 200 precision syringe pump, Chemyx, Stafford TX, USA) to perfuse 350 μL transfection reagent suspension over 1 min on top of an equal volume of Krebs solution (already in the measuring chamber). In a typical setup, this is achieved by perfusing the test solutions via a Peltier heating element. Since this arrangement does not allow attaching a Peltier element to the syringe pump, a heated plate holder mounted on the microscope table had to be implemented to allow for a constant heating of the cells at a set temperature.

### Ratiometric intracellular Ca^2+^ imaging

For intracellular Ca^2+^ imaging experiments, the cells were incubated with 6 µM Fura-2 AM (Biotium, Hayward, CA, USA) for 60 min at 37 °C in a 5% CO_2_, humidity-controlled incubator. CHO-WT were co-incubated with Pluronic F-127 20% solution in DMSO (1:40 dilution, Biotium) to facilitate cell loading. Fluorescence was measured with alternating excitation at 340 and 380 nm using a monochromator-based imaging system consisting of an MT-10 illumination system (Tokyo, Japan) and Cell-M software from Olympus. All experiments were performed at 35 °C using a standard Krebs solution. Fluorescence intensities were corrected for background signal and converted to intracellular Ca^2+^ concentrations. Only the cells that responded to the positive control (AITC) at the end of the experiment were considered TRPA1-expressing cells and included in the analysis.

### Preparation of transfection reagent solutions

Suspensions of transfection reagents were prepared at different concentrations, according to manufacturer’s recommendations, as follows: three concentrations below the recommended dose obtained by serial dilutions and two concentrations above the recommended dose. As such, for Mirus *Trans*IT, working solutions were prepared at a DNA:Mirus ratio of 1:3 [16] at the following concentrations of Mirus (μL/mL): 0.2, 0.6, 2, 6, 20, 60. For Lipofectamine 3000, working solutions were prepared at a DNA:P3000:Lipofectamine ratio of 1:2:2 [17] at the following concentrations of Lipofectamine (μL/mL): 0.5, 1.5, 3, 5, 10, 25. All lipo- and polyplexes were prepared with empty GFP vector.

### Identification of [Ca^2+^]i peaks and quantification of [Ca^2+^]i transients

In order to quantify the Ca^2+^ responses to LNP preparations, we implemented an automated analysis algorithm that: 1) identifies the baselines of the traces and corrects for the drift that arises as a result of fluorophore bleaching, 2) identifies the peaks in the traces using a dual threshold (a dynamic threshold based on the noise level of the recording and a static value of 50 nM for CHO cells and 25 nM for DRG neurons, 3) for each peak identified, it records the time of occurrence and the amplitude above the trace baseline level, and 4) for each cell, it records the number of peaks that meet the threshold criteria. Firstly, a second-order polynomial function is fitted at the bottom of the recorded traces in order to “flatten” the recordings and correct for any unwanted drift. A description of the algorithm used in our analysis (the modified polynomial algorithm) can be found here [18]. Afterwards, the corrected traces are further used to detect and quantify the Ca^2+^ transients. In short, the peak detection algorithm used for the identification of Ca^2+^ transients finds local maxima by comparing each point-value to the values of the neighboring data points within a window of desired width. If the prominence of the peak relative to its surrounding baseline exceeds a given dynamic threshold (based on the noise level of the recording), the peak is considered for further analysis. If the time distance between two consecutive peaks is smaller than a given value, the second peak is not taken into consideration. A description of the peak detection algorithm used can be found here [19,20]. The adapted Pyhton script used for analysis is freely available on GitHub [21].

### Isolation and culture of dorsal root ganglion (DRG) neurons

Primary cultures of DRG neurons were obtained from C67BL/6 wild-type mice (RRID:IMSR_JAX:000664) (12 weeks old, mixed sexes) following a protocol similar to the one described here [22]. In short, animals were euthanized in a CO_2_ chamber right before the dissection. Incisions were made under both scapulae and the neck to reach the spine. The spine was extracted by detaching it from the underlying tissue. After extraction, the spine was cleaned from any remaining muscle, tissue, and excess bone fragments. The cleaned spine was bisected, and the spinal cord was carefully removed with forceps to avoid disturbing the dural sheath membrane covering the DRGs. Bisected halves were stored on ice in a Petri dish containing PBS with penicillin and streptomycin (3%). The dural sheath covering the DRGs was removed, and the DRGs were carefully extracted using the forceps. Extracted DRGs were temporarily kept on ice and cleaned by removing the attached axons. Finally, the cleaned DRGs were stored in a 1.5 ml Eppendorf tube with basal medium (BM) (90% neurobasal medium (NBM), 10% fetal calf serum). The dissociation of DRG neurons involved two steps: chemical dissociation using 2.5 mg/mL dispase and 2 mg/mL collagenase (incubation at 37°C for one hour), followed by mechanical dissociation (via 22G and 26G syringe needles). This cell suspension was carefully layered on top of 3 ml of 16% bovine serum albumin (BSA) solution and centrifuged for 6 min at 500 x g to filter out the debris. The supernatant was removed, and the pellet was resuspended in complete medium (CM) (NBM, 0.01% GlutaMax, 0.04% B27 Supplement, 0.1% NT4, 0.01% Pen/Strep, 0.1% Glial cell-line Derived Neurotrophic Factor (GDNF)). 50 μl of the cell suspension was pipetted on each coverslip, and the coverslips were incubated for 30 min at 37°C. Finally, 2 ml of CM was added to each well. The culture has been incubated overnight in a 37°C incubator with 5% CO_2_. All phosphate-buffered saline solution (PBS) used during the culture preparation was depleted in Ca^2+^ and Mg^2+^. DRG neurons have been seeded on 13 mm coverslips, previously coated with poly-L-lysine and laminin. These experiments were approved by the KU Leuven Ethical Committee Laboratory Animals under project number (CMM-184/2021).

### Statistical analysis

Each experiment has been performed on at least 3 independent coverslips. The number of data points per condition is indicated in the caption of each figure. The statistical tests were performed using GraphPad Prism 10 software (chi-square test for proportions, Mann-Whitney test for independent skewed distributions). The definitive outliers have been identified and removed using the *Identify outliers* function of GraphPad Prism (ROUT method, Q = 0.1%). The number of Ca^2+^ transients per cell are represented as mean ± SEM, while the median amplitude of the transients is represented as box plots, where the middle line corresponds to the median value, the limits of the box correspond to the Q1 and Q3 quartiles, and the whiskers represent the 10% -90% confidence intervals. In the figures, the asterisks indicate the level of statistically significant differences (*, p < 0.05 and **, p < 0.01).

### Reagents

The following solutions have been used throughout the measurements: AITC 100 μM (Sigma-Aldrich), HC-030031 100 μM (Sigma-Aldrich), ruthenium red (RR) 10 μM (Sigma-Aldrich), pregnenolone sulfate 50 μM (Sigma-Aldrich), capsaicin 1 μM (Sigma-Aldrich), KCl 50 mM (Sigma-Aldrich). All measurements were performed in standard Krebs solution containing in mM: 150 NaCl, 6 KCl, 1.5 CaCl_2_ x 2H_2_O, 1 MgCl_2_ x 6H_2_O, 10 glucose, 5 HEPES buffer, 5 MES buffer (pH adjusted using NaOH). For the measurements performed in the absence of extracellular Ca^2+^, the Krebs solution contains the molar equivalent of CaCl_2_ in EGTA (pH adjusted using KCl).

## Results

### CHO-mTRPA1 present basal Ca^2+^ activity

We used Fura2-based intracellular Ca^2+^ imaging to assess the acute effects of lipo- and polyplexes preparations on TRPA1-expressing cells. To replicate more closely the *in vivo* scenario of slow particle diffusion through the tissue we applied the test solutions via a custom-made slow injection system (see Materials and methods), followed by a period of no perfusion to allow the undisturbed interaction of the samples with the target cells. Then, the regular gravity-driven perfusion system was turned on to wash out the cells and to apply the channel agonist (100 μM AITC), which served to identify the TRPA1-expressing cells. We performed the experiments in a thermo-regulated chamber to ensure that the suspected particle/membrane/TRPA1 interactions took place at a physiological temperature of 35 °C.

CHO-mTRPA1 cells displayed small spontaneous Ca^2+^ transients in resting conditions (i.e. no perfusion), presented as small undulatory Ca^2+^ activity. These transients were analyzed using a customized automatic algorithm that yielded the percentage of cells displaying at least one Ca^2+^ transient, the average number of Ca^2+^ transients per responding cell, and the median amplitude of Ca^2+^ transients in responding cells (see Materials and methods). Similar Ca^2+^ transient activity was observed upon injection of control Krebs solution with the syringe pump, indicating that this maneuver did not induce significant mechanical activation due to shear stress. The Ca^2+^ fluctuations recorded under basal conditions are TRPA1-mediated, as they are fully blocked by the specific channel inhibitor HC-030031 and do not occur in CHO-WT cells. Injection of AITC (10 µM) induced responses in CHO-mTRPA1 cells showing a steep ascending slope followed by a slower decay due to channel desensitization, similar to those produced by the classical continuous perfusion. This indicated that the modified setup was appropriate to assess TRPA1 responses.

As TRPA1 is a mechano- and temperature-sensitive ion channel, we expected the basal activity of CHO-mTRPA1 cells to depend on the chamber temperature, as well as on the shear stress conditions. In order to assess the involvement of these two factors to the basal Ca^2+^ measurements, we exposed CHO-mTRPA1 to four different conditions: constant perfusion of Krebs at room temperature and 35 °C, and still conditions (no liquid perfusion) at room temperature and 35 °C. As expected, the basal activity depended greatly on the measuring conditions, and was most pronounced at the physiological conditions used further in the experiments (still conditions at 35 °C) (Figure 1).

**Figure 1.**
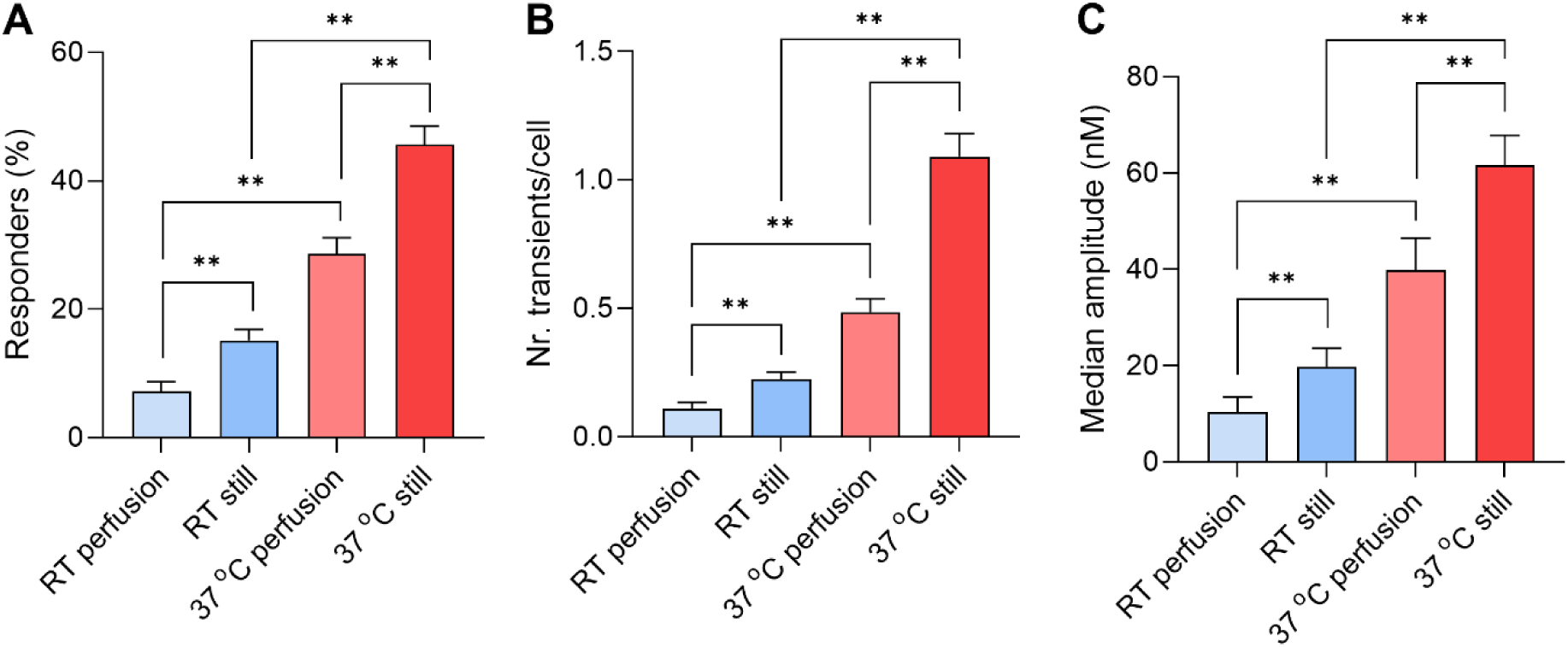
**A-C** Quantification of basal activity under different temperature and shear stress conditions in CHO-mTRPA1 cells: percentage of responding cells (A), number of Ca^2+^ transients in responding cells (B) and median amplitude of Ca^2+^ transients in responding cells (C).

### Lipofectamine 3000 triggers cytosolic Ca^2+^ transients in CHO-mTRPA1 cells

We used the microinjection system to test the effects of Lipofectamine 3000-based preparations of cationic liposomes at different concentrations (0.5, 1.5, 3, 5, 10, 25 μL/mL, denoted Lipo 0.5, Lipo 1.5, etc.) on CHO-mTRPA1 cells. Of note, Lipofectamine 3000 solutions were prepared using GFP empty vector according to manufacturer’s recommended ratios (DNA:P3000:Lipofectamine 1:2:2). These preparations and the control solution (Krebs) were characterized for size and concentration using a NTA tracker. The properties of the cationic liposome-based solutions are presented in Table 1.

**Table 1.**
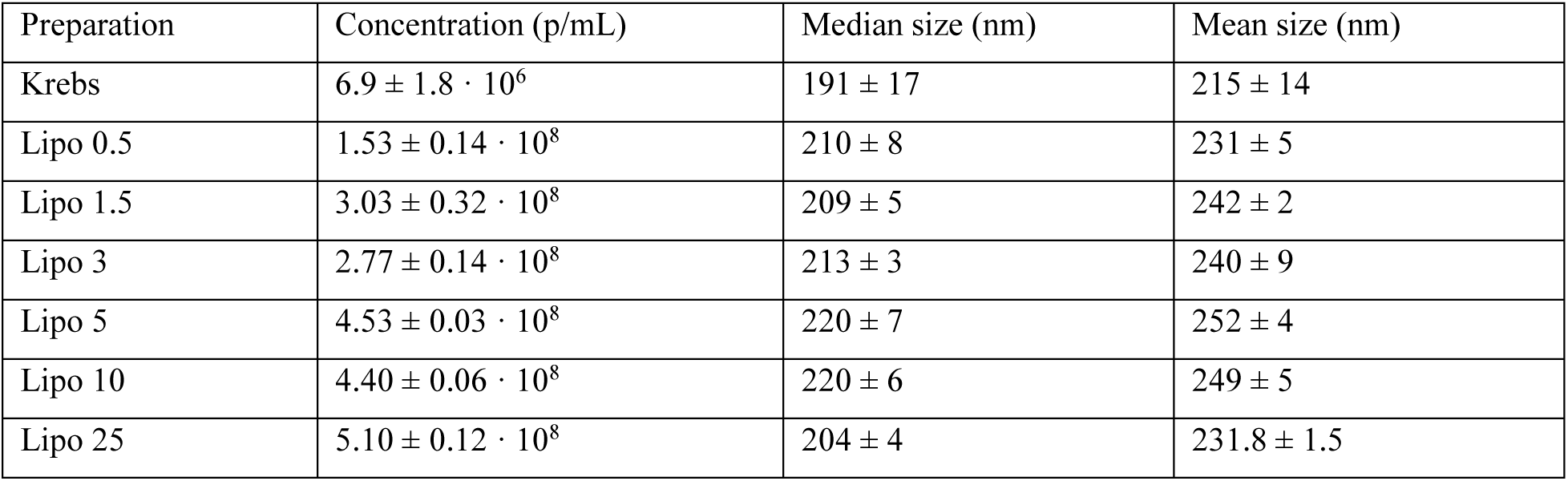
NTA characterization of Lipofectamine 3000 cationic liposomes and control solution used in the measurements.

| Preparation | Concentration (p/mL) | Median size (nm) | Mean size (nm) |
| --- | --- | --- | --- |
| Krebs | $6.9 \pm 1.8 \cdot 10^6$ | $191 \pm 17$ | $215 \pm 14$ |
| Lipo 0.5 | $1.53 \pm 0.14 \cdot 10^8$ | $210 \pm 8$ | $231 \pm 5$ |
| Lipo 1.5 | $3.03 \pm 0.32 \cdot 10^8$ | $209 \pm 5$ | $242 \pm 2$ |
| Lipo 3 | $2.77 \pm 0.14 \cdot 10^8$ | $213 \pm 3$ | $240 \pm 9$ |
| Lipo 5 | $4.53 \pm 0.03 \cdot 10^8$ | $220 \pm 7$ | $252 \pm 4$ |
| Lipo 10 | $4.40 \pm 0.06 \cdot 10^8$ | $220 \pm 6$ | $249 \pm 5$ |
| Lipo 25 | $5.10 \pm 0.12 \cdot 10^8$ | $204 \pm 4$ | $231.8 \pm 1.5$ |

Application of Lipofectamine 3000 triggered Ca^2+^ transients in the cells. These responses showed an unusual profile, compared to the typical shape of the responses to AITC. They appeared as one or multiple Ca^2+^ transients of different amplitudes throughout the application time (*Figure 2A,B*). This activation pattern may be a result of the stochastic interactions between the diffusing lipid particles and the plasma membrane of target cells.

**Figure 2.**
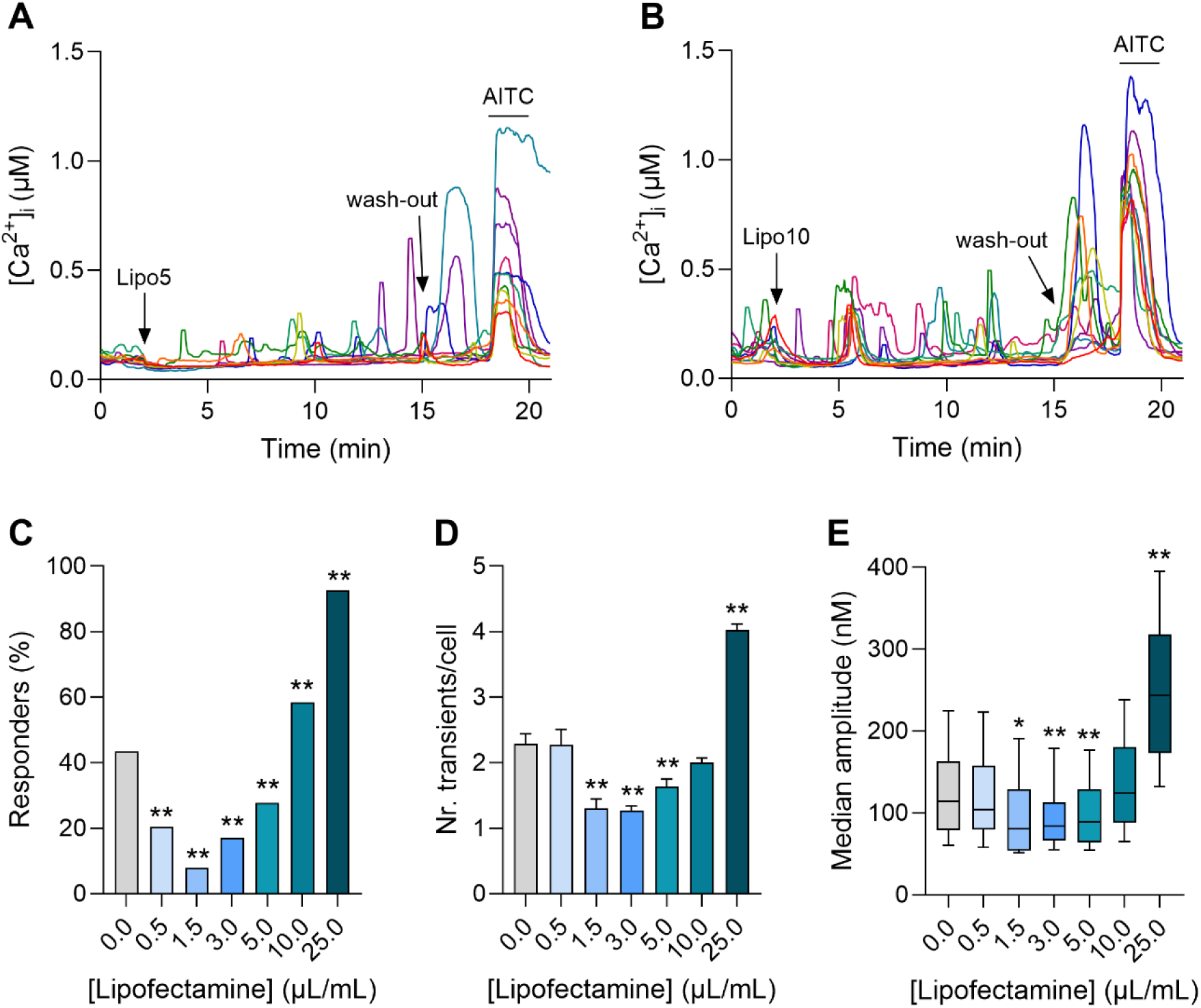
**A** Representative traces of Ca^2+^ responses in CHO-mTRPA1 cells upon injection of Lipofectamine 3000 (5 μL/mL). **B** Representative traces of Ca^2+^ responses in CHO-mTRPA1 cells upon injection of Lipofectamine 3000 (10 μL/mL). **C-D** Quantification of Ca^2+^ responses in CHO-mTRPA1 cells upon the injection of different concentrations of Lipofectamine 3000: (C) percentage of responding cells (n = 184, 264, 254, 299, 249, 536, 301 for Lipofectamine 0, 0.5, 1.5, 3, 5, 10 and 25 μL/mL), (D) number of Ca^2+^ transients in responding cells and (E) median amplitude of Ca^2+^ transients in responding cells (n = 80, 54, 20, 51, 69, 314, 279 for Lipofectamine 0, 0.5, 1.5, 3, 5, 10 and 25 μL/mL). Asterisks correspond to the comparison between each concentration of Lipofectamine 3000 and the control condition (i.e. Krebs, grey bar).

Using the customized automatic algorithm for the identification and quantification of Ca^2+^ transients, we have characterized the responses to Lipofectamine 3000 preparations in terms of 1) percentage of cells that responded with at least one Ca^2+^ transient during the time window of liposome application, 2) average number of Ca^2+^ transients per responding cell, and 3) median amplitude of Ca^2+^ transients within the subpopulation of responding cells. Interestingly, Lipofectamine 3000 preparations seem to have a dual effect on CHO-mTRPA1 cells. On one hand, Lipofectamine 3000 has an inhibitory effect on the TRPA1-dependent basal Ca^2+^ activity, which is already visible at the lowest concentrations. On the other hand, higher concentrations of cationic liposomes induce Ca^2+^ transients in CHO-mTRPA1 cells, leading to a J-shaped dose response curve. Of note, the net Ca^2+^ activity in the cells challenged with the typical dose recommended for a successful transfection (Lipo 5) does not exceed the one measured under basal conditions (injection of Krebs) (*Figure 2C-E*).

### Lipofectamine 3000 preparations inhibit the basal Ca^2+^ activity mediated by TRPA1

As mentioned in the previous section, low doses of Lipofectamine 3000 seem to inhibit the basal activity of CHO-mTRPA1 cells (*Figure 2C-E*). Based on the parameter tested, the strongest inhibition is visible for Lipo 1.5 and Lipo 3. As the basal activity of CHO-mTRPA1 cells is TRPA1-mediated, we tested the hypothesis that Lipofectamine 3000 preparations induce TRPA1 channel inhibition.

A first indication of TRPA1 inhibition came from comparing the average Ca^2+^ responses to AITC at the end of the protocol (that is, after the wash-out of liposome preparations). As can be observed in *Figure 3A*, the AITC responses after the application of Lipo 3 are smaller in amplitude and delayed compared to the ones measured after the application of Krebs (values normalized to AITC responses measured after the application of Krebs). This effect is dose-dependent and it peaks at concentration Lipo 3, after which it reaches a plateau. A second indication of TRPA1 inhibition by Lipofectamine 3000 is the rebound response during the wash-out phase (minute 15 to 18) (*Figure 2A,B*), which is characteristic to TRPA1 channel inhibitors. Of note, the rebound responses did not correlate with the concentration of Lipofectamine 3000 applied.

**Figure 3.**
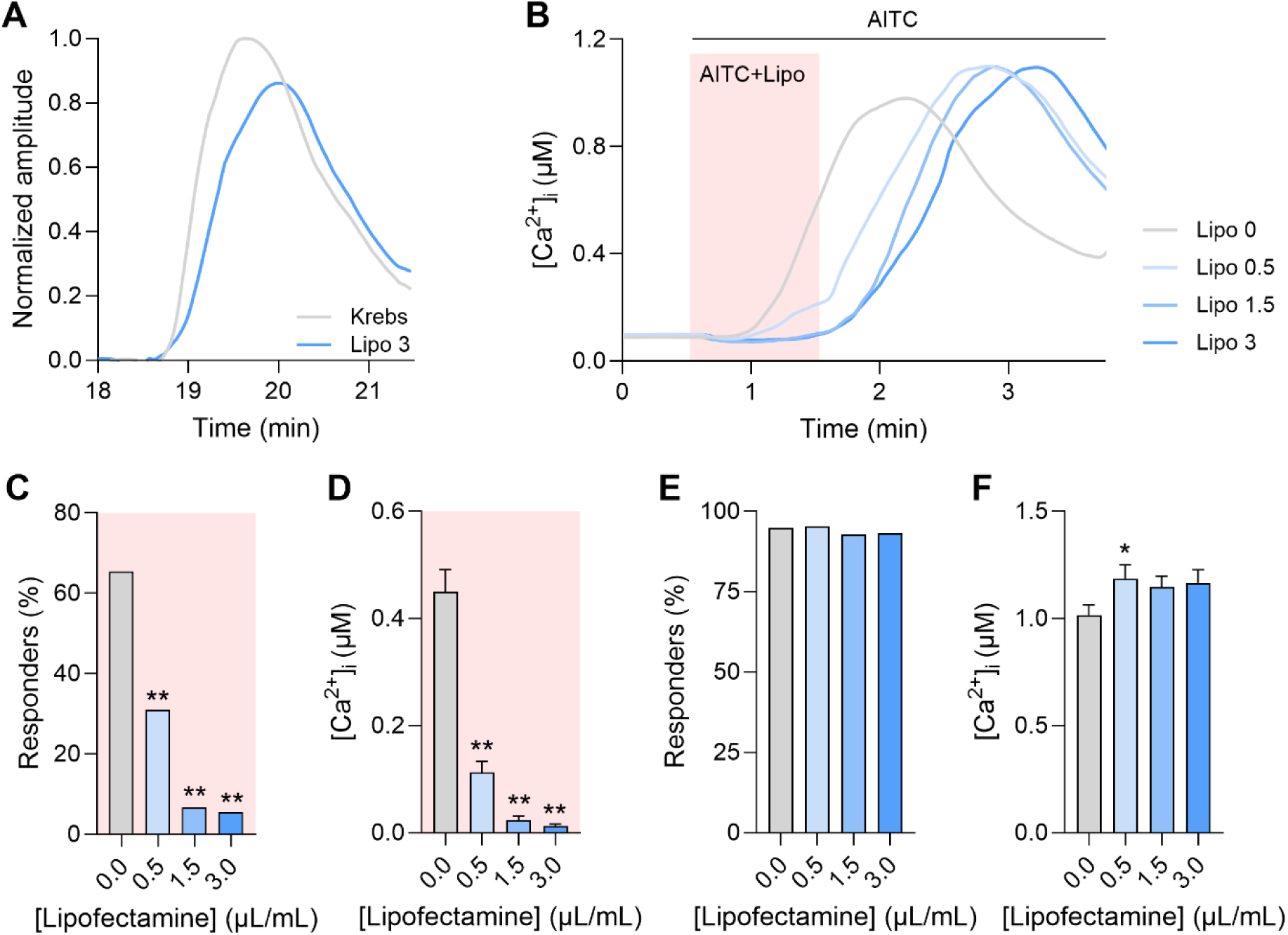
TRPA1 inhibition by Lipofectamine 3000. **A** Normalized amplitude of average AITC responses at the end of the protocol (after Krebs, respectively liposome wash-out). **B** Average responses of AITC and Lipofectamine co-application, followed by Lipofectamine wash-out. **C,D** Quantified Ca^2+^ responses corresponding to the co-application of AITC and Lipofectamine (minute 0.5 to 1.5 in panel B): (C) percentage of responding cells and (D) average Ca^2+^ amplitude of the whole population (n = 191, 129, 210, 145 for Lipofectamine 0, 0.5, 1.5 and 3 μL/mL). **E,F** Quantified Ca^2+^ responses corresponding to Lipofectamine wash-out (minute 1.5 –3.5 in panel B): (E) percentage of responding cells and (F) average Ca^2+^ amplitude of the whole population (n = 191, 129, 210, 145 for Lipofectamine 0, 0.5, 1.5 and 3 μL/mL). The asterisks correspond to the comparison between each concentration of Lipofectamine and the vehicle solution Krebs (i.e. Lipo 0).

To better assess the inhibitory effects of Lipofectamine 3000 on TRPA1, we performed another set of measurements, in which AITC was applied either on its own, or in the presence of varying concentrations of Lipofectamine 3000 (Lipo 0.5, Lipo 1.5 and Lipo 3) (*Figure 3B*). After 1.5 min, liposomes were washed out and AITC was applied on its own. For these measurements, we have characterized the responses to both time intervals (AITC + Lipo and AITC alone) in terms of percentage of responding cells and average Ca^2+^ amplitude in the whole population (*Figure 3C,D*). As can be seen in the quantified responses, Lipofectamine 3000 preparations have a dose-dependent inhibitory effect on TRPA1, with virtually full inhibition already achieved by Lipo 1.5. However, after liposome wash-out, AITC responses immediately returned to normal (*Figure 3E,F*). This confirms that Lipofectamine 3000 preparations act as a potent TRPA1 channel inhibitor, and concentrations below the manufacturer’s recommended dose are effective in blocking the AITC-induced Ca^2+^ responses.

### The Lipofectamine 3000-induced Ca^2+^ transients are partly mediated by TRPA1

To test the hypothesis that TRPA1 is involved in the liposome-induced generation of Ca^2+^ responses, we have applied the Lipofectamine 3000 preparations in the presence of the specific TRPA1 inhibitor HC-030031 and the non-specific Ca^2+^ channel blocker ruthenium red (RR). For this, we focused on the effects of the manufacturer’s recommended dose (Lipo 5) and the next higher concentration (Lipo 10). Representative traces of Ca^2+^ responses elicited by the co-application of Lipo 10 and the two TRPA1 blockers can be observed in *Figure 4A,B*. Interestingly, the specific TRPA1 inhibitor only reduced the percentage of responding cells activated by Lipo 10, suggesting a partial involvement of TRPA1 in the generation of the Ca^2+^ transients. While the non-specific Ca^2+^ channel blocker RR successfully reduced the Ca^2+^ responses induced by Lipo 10 as described by all tested parameters, it only had an effect on the percentage of responding cells activated by the typical working concentration (Lipo 5). For the quantified responses in the presence of the channel blockers please refer to *Figure 4C-E*. The differential inhibitory effect of HC-030031 on the two liposome concentrations tested may suggest that the involvement of TRPA1 in Lipofectamine 3000-induced Ca^2+^ responses is dose-dependent, and this effect only becomes visible at higher doses (starting from Lipo 10). Another important aspect to consider is that the Lipofectamine 3000 preparations are themselves potent TRPA1 inhibitors, as proven in the previous section. As Lipo 3 is already effective in almost fully blocking the TRPA1 channel, the specific inhibitor HC-030031 is not able to exert an additional inhibitory effect (since the channel is already fully blocked). The non-specific TRPA1 blocker RR is able to induce further inhibitory effects (compared to HC-030031), suggesting an additional, TRPA1-independent pathway involved in the generation of Lipofectamine 3000-induced Ca^2+^ transients. However, RR does not fully abolish the Ca^2+^ responses, suggesting that they do not solely arise as Ca^2+^ influx from the extracellular medium. In fact, in measurements performed in the absence of extracellular Ca^2+^, Lipo 5 and Lipo 10 were able to induce Ca^2+^ transients in approximately 70% of the cells . This points towards a third mechanism involved in liposome-induced Ca^2+^ responses in CHO-mTRPA1, namely Ca^2+^ mobilization from intracellular stores. In conclusion, at least three mechanisms are involved in Lipofectamine 3000-induced Ca^2+^ transients in CHO-mTRPA1 cells: TRPA1-dependent Ca^2+^ influx, TRPA1-independent Ca^2+^ influx (via an unknown mechanism) and Ca^2+^ mobilization from internal stores.

**Figure 4.**
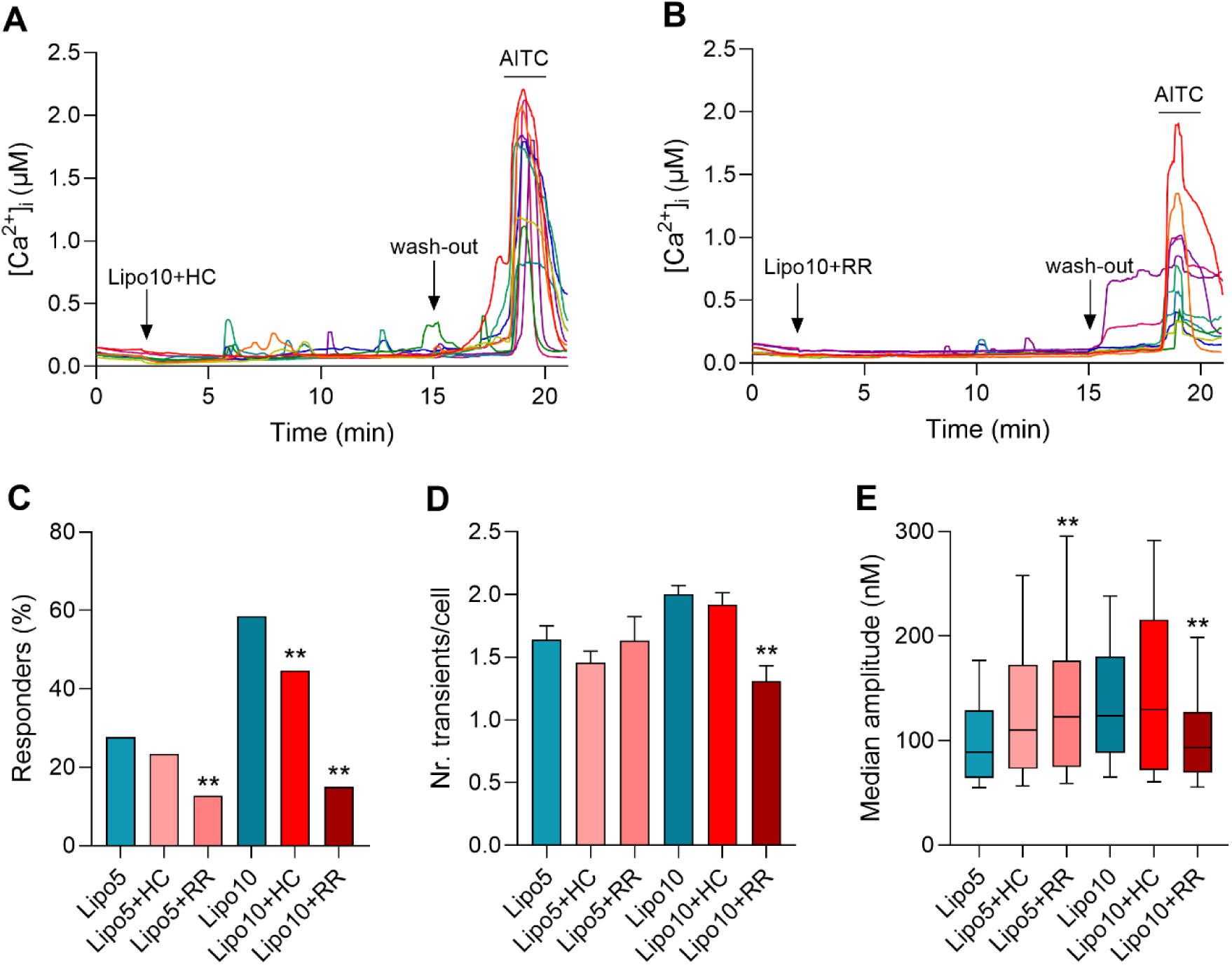
**A** Representative traces of Ca^2+^ responses elicited by Lipo 10 in the presence of the specific TRPA1 blocker HC-030031. **B** Representative traces of Ca^2+^ responses elicited by Lipo 10 in the presence of the non-specific Ca^2+^ channel blocker RR. **C-E** Quantification of Ca^2+^ transients elicited in the absence and presence of TRPA1 blockers for two different concentrations of Lipofectamine (Lipo 5 and Lipo 10): (C) percentage of responding cells (n = 249, 401, 213, 536, 325, 194 for Lipo5, Lipo5+HC, Lipo5+RR, Lipo10, Lipo10+HC and Lipo10+RR), (D) average number of Ca^2+^ transients per cell in responding cells and (E) median amplitude of Ca^2+^ transients in responding cells (n = 69, 94, 27, 314, 145, 29 for Lipo5, Lipo5+HC, Lipo5+RR, Lipo10, Lipo10+HC and Lipo10+RR). The asterisks correspond to the comparison between the responses in the presence of the blockers and in the absence of the blockers for each concentration of Lipofectamine.

### Lipofectamine 3000 preparations trigger cytosolic Ca^2+^ transients in CHO-WT cells

Since the channel blockers do not fully inhibit the responses induced by Lipofectamine 3000 in CHO-mTRPA1 cells, we decided to explore the possibility that these preparations might have other targets. For this, we performed the same kind of measurements, but in CHO-WT cells. Firstly, the CHO-WT do not show basal activity upon the injection of Krebs (*Figure 5A*). The application of cationic liposome preparations resulted in the same type of transient and variable Ca^2+^ responses in CHO-WT, as observed in CHO-mTRPA1 cells (*Figure 5B*). These responses are dose-dependent and, unlike the case of CHO-mTRPA1 cells, the dose response curve is linear, rather than J-shaped, as CHO-WT cells do not present basal activity. The dose response curves measured in CHO-WT cells are shown in *Figure 5C-E*, together with CHO-mTRPA1 curves for reference. Apart from the highest concentration tested (i.e. Lipo 25), CHO-WT cells respond at least as strongly as the CHO-mTRPA1 counterpart. It is possible that the inhibitory action of Lipofectamine on the TRPA1 channel prevents CHO-mTRPA1 cells from responding stronger to these preparations. Another possible explanation for the relatively high responses in CHO-WT cells may be related to the use of Pluronic F-127 during the Fura2-AM incubation time. Pluronic is a surfactant used to increase the efficiency of Fura2-AM loading in CHO-WT cells that, unlike CHO-mTRPA1 cells do not uptake the fluorophore on their own. The interaction between the surfactant and plasma membrane may confer CHO-WT cells different properties compared to the naïve state (i.e. altered plasma membrane fluidity).

**Figure 5.**
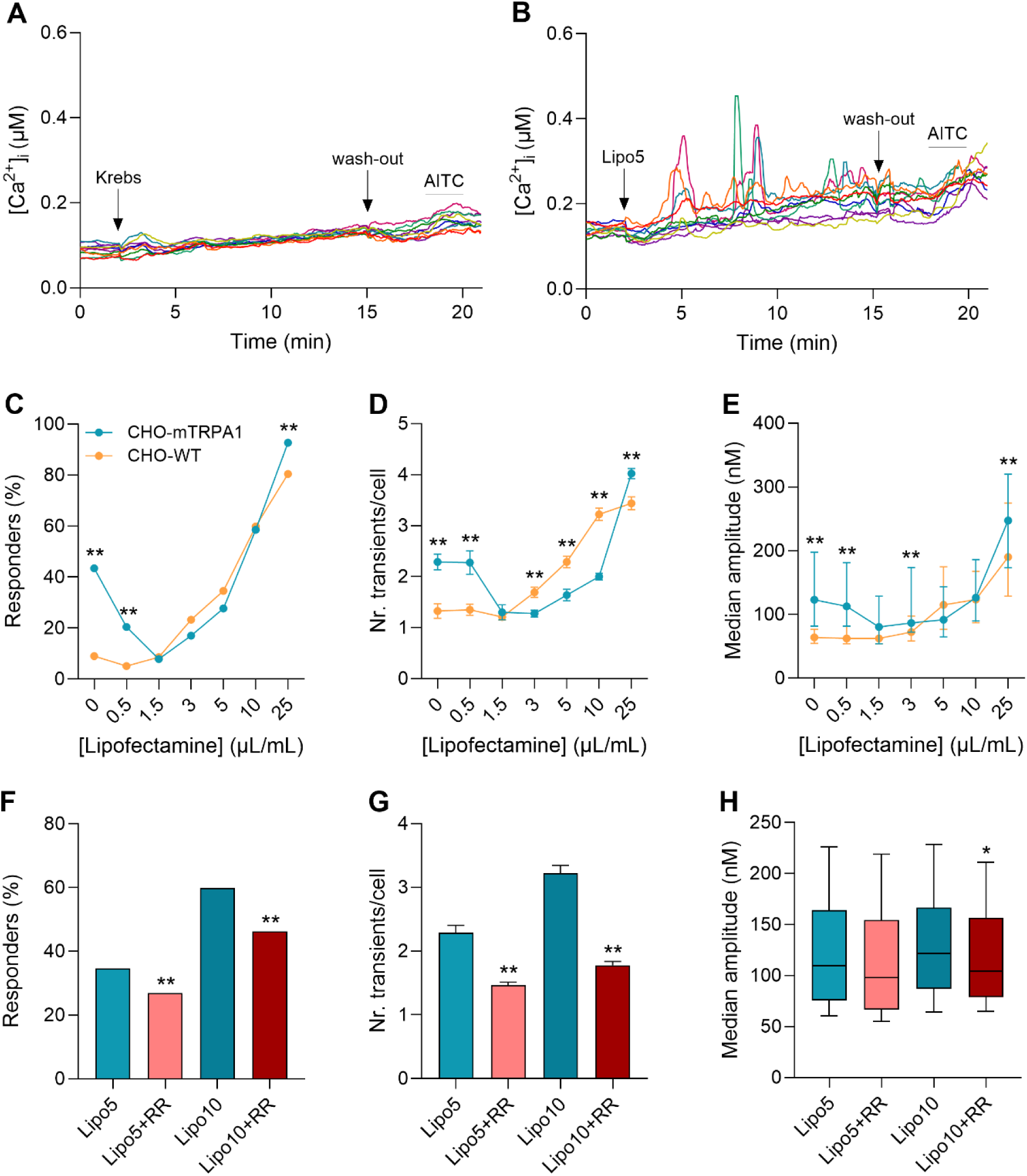
Lipofectamine 3000 preparations induce Ca^2+^ transients in CHO-WT cells. **A** Representative traces of Ca^2+^ responses in CHO-WT cells upon injection of Krebs. **B** Representative traces of Ca^2+^ responses in CHO-WT cells upon injection of Lipofectamine 3000 (5 μL/mL). **C-E** Dose-response curves measured in CHO-WT (orange) and CHO-mTRPA1 (teal) cells: (C) percentage of responding cells (n = 644, 786, 763, 515, 479, 526, 429 for Lipofectamine 0, 0.5, 1.5, 3, 5, 10 and 25 μL/mL in CHO-WT cells), (D) average number of Ca^2+^ transients per cell in responding cells and (E) median amplitude of Ca^2+^ responses in responding cells (n = 58, 40, 66, 120, 166, 315, 345 for Lipofectamine 0, 0.5, 1.5, 3, 5, 10 and 25 μL/mL in CHO-WT cells). Asterisks correspond to the comparison between different doses of Lipofectamine and the vehicle solution Krebs (i.e. Lipo 0). **F-H** Quantification of Ca^2+^ transients elicited by Lipofectamine in the absence and presence of the non-specific Ca^2+^ channel blocker RR: (F) percentage of responding cells (n = 479, 1157, 526, 666 for Lipo5, Lipo5+RR, Lipo10 and Lipo10+RR), (G) average number of Ca^2+^ transients per cell in responding cells and (H) median amplitude of Ca^2+^ transients in responding cells (n = 166, 311, 315, 308 for Lipo5, Lipo5+RR, Lipo10 and Lipo10+RR). Asterisks correspond to the comparison between the responses in the presence and absence of RR, for each concentration of Lipofectamine.

To test whether the increase in cytosolic Ca^2+^ is a result of influx from the extracellular medium in CHO-WT cells, we used the non-specific Ca^2+^ channel blocker RR. As illustrated in *Figure 5F-H*, RR successfully reduced the responses to Lipofectamine preparations, pointing towards Ca^2+^ influx from the extracellular medium. However, like in the case of CHO-mTRPA1 cells, RR was not able to abolish Lipofectamine-induced Ca^2+^ transients. When applied in the absence of extracellular Ca^2+^, Lipo 5 and Lipo 10 triggered Ca^2+^ responses in 9%, respectively 47% of the population. Similar to CHO-mTRPA1 cells, multiple pathways are involved in the generation of Lipofectamine preparations-induce Ca^2+^ responses in CHO-WT cells: on one hand, Ca^2+^ influx from the extracellular medium via a yet-to-be-identified mechanism and, on the other hand, Ca^2+^ mobilization from internal stores.

### Lipofectamine 3000 components have differential effects in target cells

As Lipofectamine 3000 is a transfection reagent composed of two separate reagents that work together to enhance the transfection efficiency (i.e. Lipofectamine and P3000), we wanted to test the effects of each of them on target cells. Because Lipofectamine 3000 preparations seem to have a dual effect on CHO-mTRPA1 cells (simultaneous inhibition and activation), we chose to investigate the effects of 10 μL/mL of each reagent (no nucleic acids added) in CHO-WT cells, where only the stimulatory effect is visible. We chose this concentration because it induces very clear and relatively large responses. As can be seen in the example traces displayed in *Figure 6A,B*, the two reagents of Lipofectamine 3000 have differential effects in CHO-WT cells. While Lipofectamine on its own only induces minimal cytosolic Ca^2+^ responses, comparable to the ones induced by the control vehicle Krebs, the enhancer P3000 triggered significantly more Ca^2+^ transients. Moreover, the mixture of the two reagents results in enhanced Ca^2+^ responses (*Figure 6C-D*).

**Figure 6.**
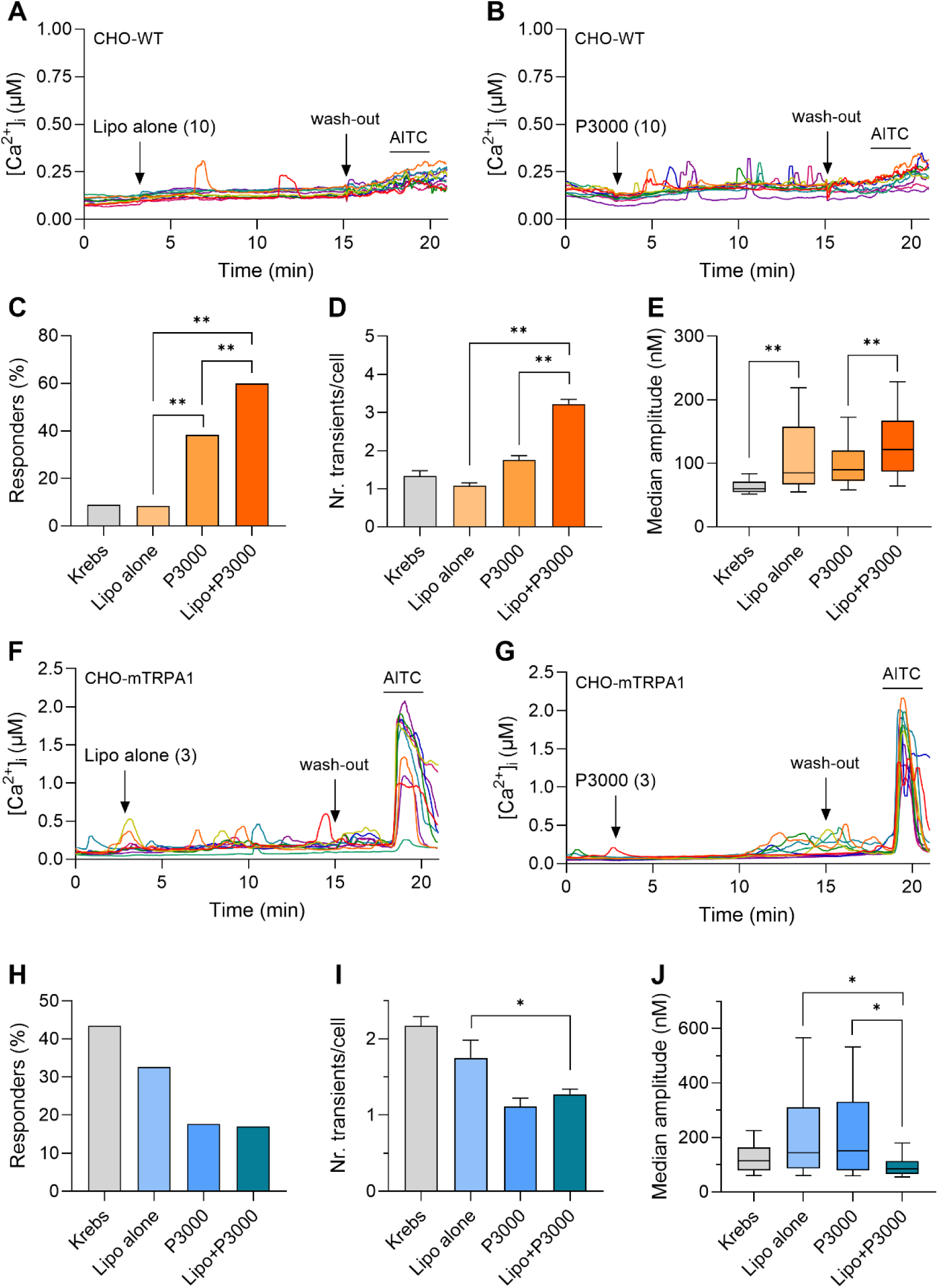
**A** Representative traces of Ca^2+^ responses induced by Lipofectamine (10 μL/mL) alone in CHO-WT cells. **B** Representative traces of Ca^2+^ responses induced by the P3000 enhancer (10 μL/mL) in CHO-WT cells. **C-E** Quantification of Ca^2+^ transients induced by Krebs, Lipo 10 and the two reagents individually (at 10 μL/mL): (C) percentage of responding cells (n = 644, 154, 224, 526 for Krebs, Lipo alone, P3000 and Lipo+P3000), (D) average number of Ca^2+^ transients per cell in responding cells, (E) median amplitude of Ca^2+^ transients in responding cells (n = 58, 13, 86, 315 for Krebs, Lipo alone, P3000 and Lipo+P3000). **F** Representative traces of Ca^2+^ responses induced by Lipofectamine (3 μL/mL) alone in CHO-mTRPA1 cells. **G** Representative traces of Ca^2+^ responses induced by the P3000 enhancer (3 μL/mL) in CHO-mTRPA1 cells. **H-J** Quantification of Ca^2+^ transients induced by Krebs, Lipo 3 and the two reagents individually (at 3 μL/mL): (H) percentage of responding neurons (n = 184, 49, 51, 299 for Krebs, Lipo alone, P3000 and Lipo+P3000), (I) average number of Ca^2+^ transients per cell in responding cells, (J) median amplitude of Ca^2+^ transients in responding cells (n = 99, 16, 9, 51 for Krebs, Lipo alone, P3000 and Lipo+P3000).

Next, we assessed the inhibitory effects of the two components of Lipofectamine 3000 on the basal activity of CHO-mTRPA1 cells (*Figure 6F-J*). For this, we used 3 μL/mL of each reagent, as at this concentration, Lipo 3 caused the maximal inhibitory effect on TRPA1 (*Figure 3B-D*). While Lipofectamine on its own does not significantly affect the basal Ca^2+^ activity in the cells, P3000 causes a clear inhibition of the TRPA1-dependent basal activity. The strongest inhibitory effect is triggered by the combination of both reagents (*Figure 6H-J*).

These experiments suggest that Lipofectamine on its own does not cause significant alteration of Ca^2+^ homeostasis in target cells, while the P3000 enhancer seems to be mainly responsible for both the TRPA1 inhibitory effect, as well as the generation of Ca^2+^ transients in target cells. The TRPA1-dependent stimulatory effect of Lipofectamine 3000 components in CHO-mTRPA1 cells is not easy to assess, since this effect cannot be studied independently from TRPA1 inhibition.

### Lipofectamine 3000 induces Ca^2+^ transients in DRG neurons

TRPA1 is most endogenously expressed in sensory neurons, where it acts as a nociceptive receptor. Hence, primary cultures of DRG neurons cell bodies have been used to perform Ca^2+^-imaging, in order to assess the effects of cationic liposomes. Similar to CHO cells, the application of Lipofectamine 3000 preparations induced Ca^2+^ transients in DRG neurons (*Figure 7A*). Unlike CHO-mTRPA1 cells, DRG neurons present no basal Ca^2+^ activity and more stable and flatter baselines, which allowed for lowering the detection threshold for Ca^2+^ transients from 50 nM to 25 nM. The Ca^2+^ responses are dose-dependent, although smaller in amplitude compared to CHO-WT and CHO-mTRPA1 cells (*Figure 5C-E* and *Figure 7C-E*).

**Figure 7.**
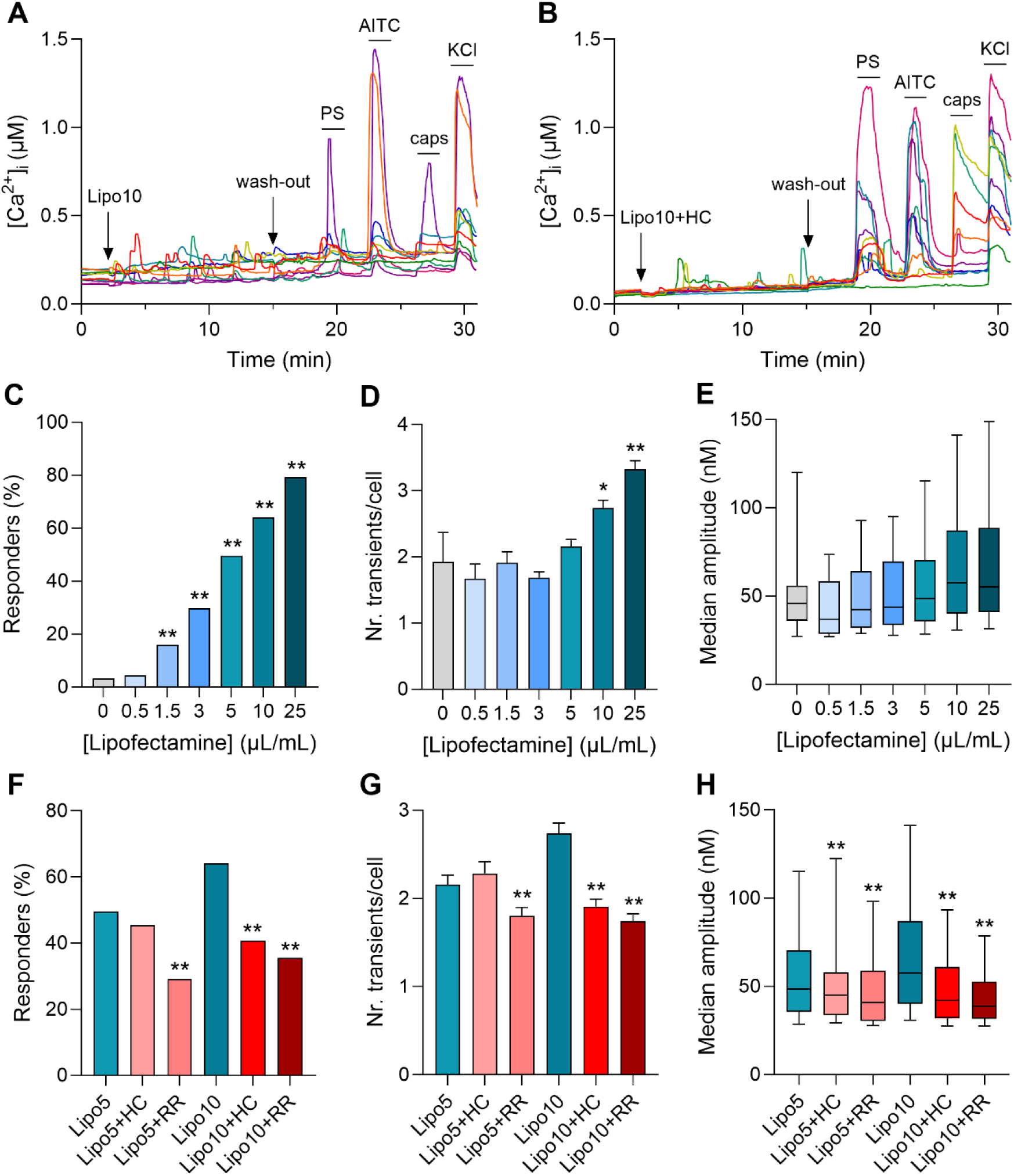
**A** Representative traces of Ca^2+^ responses induced by Lipo 10 in DRG neurons. **B** Representative traces of Ca^2+^ responses induced by Lipo 10 in the presence of the specific TRPA1 blocker HC-030031. **C-E** Quantification of dose-dependent Ca^2+^ responses induced by increasing concentrations of Lipofectamine 3000: (C) percentage of responding neurons (n = 402, 603, 483, 456, 351, 318, 322 for Lipofectamine 0, 0.5, 1.5, 3, 5, 10 and 25 μL/mL), (D) average number of Ca^2+^ transients per cell in responding neurons, (E) median amplitude of Ca^2+^ transients in responding neurons (n = 13, 27, 77, 136, 174, 203, 255 for Lipofectamine 0, 0.5, 1.5, 3, 5, 10 and 25 μL/mL). Asterisks correspond to the comparison between different doses of Lipofectamine and the vehicle solution Krebs (i.e. Lipo 0). **F-H** Quantification of Ca^2+^ transients recorded in the absence and in the presence of TRPA1 blockers: (F) percentage of responding neurons (n = 351, 301, 588, 318, 558, 529 for Lipo5, Lipo5+HC, Lipo5+RR, Lipo10, Lipo10+HC and Lipo10+RR), (G) average number of Ca^2+^ transients per cell in responding neurons, (H) median amplitude of Ca^2+^ transients in responding neurons (n = 174, 137, 171, 204, 228, 188 for Lipo5, Lipo5+HC, Lipo5+RR, Lipo10, Lipo10+HC and Lipo10+RR). Asterisks correspond to the comparison between the responses in the presence and absence of HC-030031, respectively RR, for each concentration of Lipofectamine.

To test whether the Ca^2+^ transients recorded in DRG neurons are TRPA1-mediated, Lipofectamine preparations have been applied in the presence of the specific TRPA1 channel inhibitor HC-030031 and the non-specific Ca^2+^ channel blocker RR. HC-030031 significantly reduced the Ca^2+^ responses induced by Lipo 10 in the whole population of neurons across all tested parameters, while in the case of Lipo 5 it had an effect only on the median amplitude of the responses (*Figure 7F-H*). Of note, a complete reduction to baseline levels has not been achieved in any of the cases. This strongly points to a partial involvement of TRPA1 in the generation of Lipofectamine 3000-induced Ca^2+^ responses in DRG neurons, which may also be dose-dependent (i.e. the lower concentrations tested failed to contribute to the overall responses in a TRPA1-dependent manner).

As most of the Ca^2+^ signals triggered by Lipofectamine preparations have not been successfully abolished by the specific TRPA1 inhibitor HC-030031, it is possible that another ion channel also mediates Ca^2+^ influx in DRG neurons. To test this hypothesis, Lipofectamine preparations have been applied in the presence of RR. As shown in *Figure 7F-H*, RR significantly reduced Ca^2+^ transients, as measured by all the parameters. Moreover, it caused an additional reduction compared to HC-030031, suggesting additional TRPA1-independent mechanisms of Ca^2+^ influx involved in the generation of Lipofectamine 3000-induced Ca^2+^ transients in DRG neurons. In line with the observations from CHO-mTRPA1 cells, RR did not abolish the Ca^2+^ transients. This indicated that cationic liposomes may also trigger Ca^2+^ mobilization from internal stores. Indeed, when applied in the absence of extracellular Ca^2+^, Lipo 5 and Lipo 10 induced activation in 8.5%, respectively 11.6% of the total DRG neuron population. This confirms that, similar to CHO-mTRPA1 cells, Lipofectamine 3000 preparations induce Ca^2+^ responses in DRG neurons via at least three mechanisms: TRPA1-dependent Ca^2+^ influx, TRPA1-independent Ca^2+^ influx (via an unknown pathway) and Ca^2+^ mobilization from intracellular stores.

#### Mirus *Trans*IT preparations trigger Ca^2+^ transients in target cells

To assess whether the ability of chemical transfection reagents is limited to cationic liposomes, we used the microinjection system to apply a different type of chemical transfection reagent based on cationic polymers, namely Mirus *Trans*IT-293. For this, we tested multiple concentrations of Mirus, preparations centered around the manufacturer’s recommended dose of 6 μL/mL. The NTA characterization of the different preparations used in these measurements is summarized in Table 2.

**Table 2.** NTA characterization of Mirus *Trans*IT preparations.

| Preparation | Concentration (p/mL) | Median size (nm) | Mean size (nm) |
| --- | --- | --- | --- |
| Krebs | $3.83 \pm 0.26 \cdot 10^6$ | $226 \pm 16$ | $244 \pm 16$ |
| Mirus 0.6 | $4.1 \pm 0.5 \cdot 10^7$ | $219 \pm 14$ | $275 \pm 9$ |
| Mirus 2 | $4.6 \pm 0.4 \cdot 10^7$ | $194 \pm 13$ | $223 \pm 10$ |
| Mirus 6 | $1.0 \pm 0.3 \cdot 10^8$ | $197 \pm 5$ | $218 \pm 5$ |
| Mirus 20 | $1.2 \pm 0.5 \cdot 10^9$ | $206 \pm 19$ | $226 \pm 18$ |

In all cell types tested (CHO cells and DRG neurons), the application of cationic polymer preparations triggered similar Ca^2+^ transients as Lipofectamine 3000-derived cationic liposomes, described in the previous sections (*Figure 8A,H*). Interestingly, in CHO-mTRPA1 cells, Mirus *Trans*IT preparations have a bell-shaped dose response which peaks, in the case of all parameters tested, at the reference concentration of 6 μL/mL (*Figure 8B-D*). Below this concentration, Mirus preparations do not induce significant Ca^2+^ signals above the basal activity (with the exception of median Ca^2+^ transient amplitude measured for Mirus 2). Above the dose of Mirus 6, the Ca^2+^ responses return to baseline level. This suggests an inhibitory effect of high doses of Mirus *Trans*IT preparations, an effect observed for Lipofectamine 3000 as well. In contrast to Lipofectamine 3000, for which the inhibitory effect is observed in the lower range of the concentration, Mirus *Trans*IT exhibits an inhibitory effect towards the higher end of concentrations.

**Figure 8.**
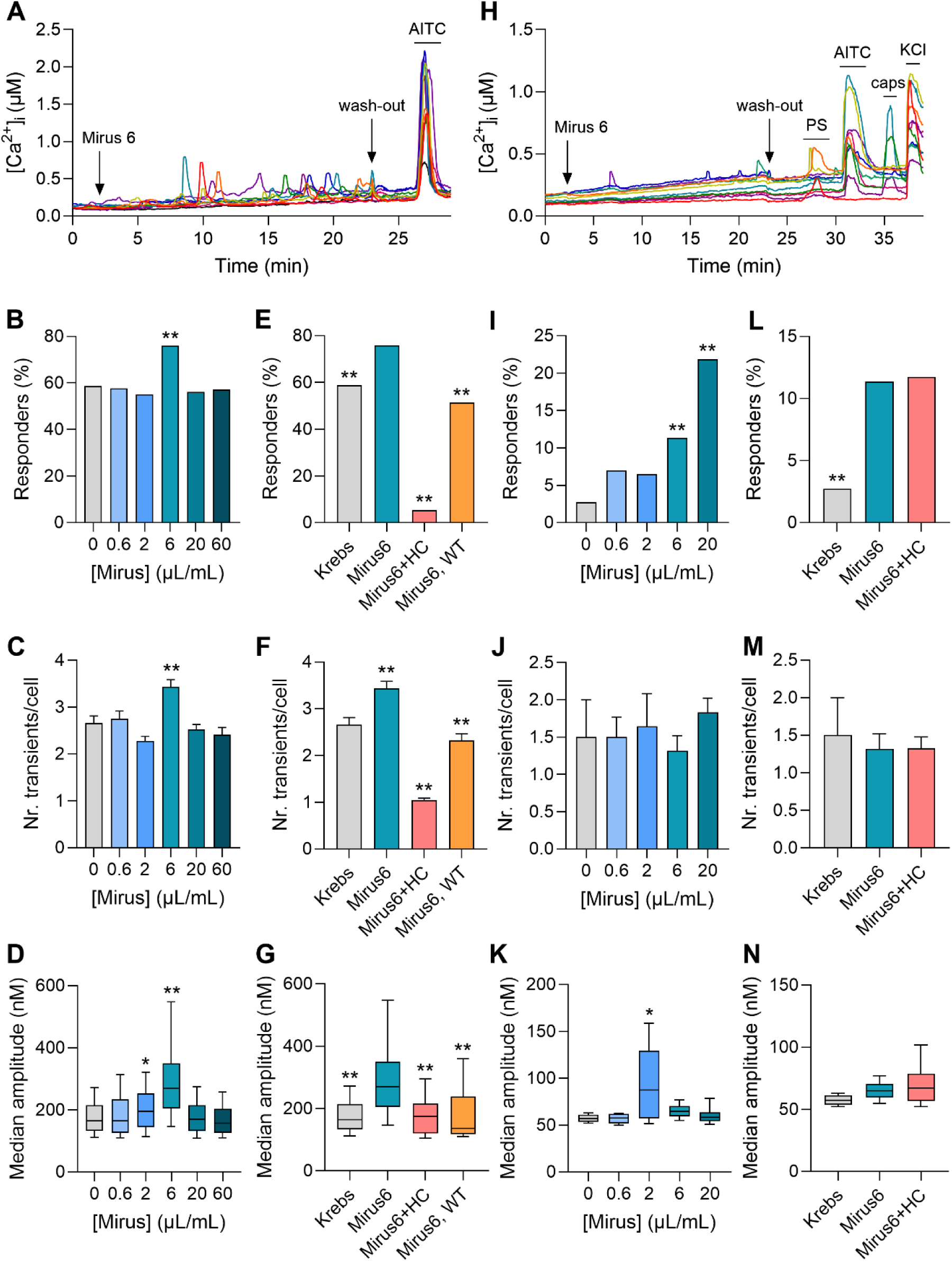
Mirus *Trans*IT-293 preparations induce Ca^2+^ signals in target cells. **A** Representative traces of Ca^2+^ transients triggered by Mirus 6 preparations in CHO-mTRPA1 cells. **B-D** Dose-response curves of Mirus preparations, as measured by different parameters of the responses: (B) percentage of responding cells (n = 172, 189, 303, 170, 340, 170 for Mirus 0, 0.6, 2, 6, 20 and 60 μL/mL), (C) number of Ca^2+^ transients per cell in responding cells, and (D) median amplitude of Ca^2+^ transients in responding cells (n = 101, 109, 167, 129, 191, 97 for Mirus 0, 0.6, 2, 6, 20 and 60 μL/mL). Asterisks correspond to the comparison between each concentration and the vehicle control solution Krebs. **E-G** Quantification of Ca^2+^ transients elicited by different negative controls (control vehicle solution Krebs in CHO-mTRPA1 cells, Mirus 6 application in the presence of the specific TRPA1 blocker HC-030031 in CHO-mTRPA1 cells, and Mirus 6 application in CHO-WT cells): (E) percentage of responding cells (n = 172, 170, 421, 232 for Krebs, Mirus6, Mirus6+HC, Mirus6, WT), (F) number of Ca^2+^ transients per cell in responding cells, and (G) median amplitude of Ca^2+^ transients in responding cells (n = 101, 129, 23, 119 for Krebs, Mirus6, Mirus6+HC, Mirus6, WT). Asterisks correspond to the comparison between each negative control condition and the responses elicited by Mirus 6 in CHO-mTRPA1 cells. **H** Representative traces of Ca^2+^ transients triggered by Mirus 6 preparations in DRG neurons. **I-K** Dose-response curves of Mirus preparations, as measured by different parameters of the responses: (I) percentage of responding cells (n = 219, 143, 215, 194, 160 for Mirus 0, 0.6, 2, 6 and 20 μL/mL), (J) number of Ca^2+^ transients per cell in responding cells, and (K) median amplitude of Ca^2+^ transients in responding cells (n = 6, 10, 14, 22, 35 for Mirus 0, 0.6, 2, 6 and 20 μL/mL). Asterisks correspond to the comparison between each concentration and the vehicle control solution Krebs. **L-N** Quantification of Ca^2+^ transients elicited by different negative controls (control vehicle solution Krebs and Mirus 6 application in the presence of the specific TRPA1 blocker HC-030031): (L) percentage of responding cells (n = 219, 194, 239 for Krebs, Mirus6 and Mirus6+HC), (M) number of Ca^2+^ transients per cell in responding cells, and (N) median amplitude of Ca^2+^ transients in responding cells (n = 6, 22, 28 for Krebs, Mirus6 and Mirus6+HC). Asterisks correspond to the comparison between each negative control condition and the responses elicited by Mirus 6 in CHO-mTRPA1 cells.

In order to assess the contribution of TRPA1 to the Mirus *Trans*IT-293-induced Ca^2+^ transients, the cationic polymer preparations have been applied in the presence of the specific TRPA1 inhibitor HC-030031. *Figure 8E-G* shows that HC-030031 drastically reduces the Ca^2+^ transients, as viewed by all tested parameters. In fact, it reduced the percentage of responding cells and the number of transients per cell well below the level of basal activity, while the median amplitude of Ca^2+^ responses in responding cells is returned to basal levels. Moreover, the application of Mirus 6 in CHO-WT cells elicits significantly lower Ca^2+^ responses compared to CHO-mTRPA1 cells. This, together with the strong inhibitory effects of HC-030031, suggests a partial implication of TRPA1 in the generation of Ca^2+^ transients evoked by Mirus *Trans*IT-293-derived cationic polymer structures.

In DRG neurons, Mirus preparations trigger far less and smaller Ca^2+^ transients compared to CHO cells (*Figure 8I-K*). Interestingly, the dose-response curves follow different shapes, depending on the parameter tested. For example, when assessing the percentage of activated neurons, the dose-response follows a monotonically increasing function (*Figure 8I,J*), while in the case of median amplitude of Ca^2+^ transients in responding cells, the dose-response is bell-shaped and it peaks of the concentration of 2 μL/mL (*Figure 8K*). These differences may be a consequence of the fact that DRG neurons are complex systems expressing a multitude of membrane receptors, including different TRP channels (as highlighted by the agonist responses at the end of the stimulation protocol). It is possible that multiple of these membrane receptors are sensitive to Mirus preparations, and the overall activation curve observed in DRG neurons is the cumulative effect on each of the receptors (some of them being activated, while some of them being inhibited by different concentrations). Nevertheless, the contribution of TRPA1 to these responses seems to be minimal in DRG neurons, as the specific TRPA1 inhibitor HC-030031 does not reduce the Ca^2+^ signals triggered by the cationic polymer preparations (*Figure 8L-N*). This strengthens the hypothesis that other molecular players may be targeted by Mirus *Trans*IT-293 as well.

## Discussion

In this study, we tested the ability of different types of nanocarriers to modulate the activity of TRPA1. For this, we used two types of commercially available transfection reagents based that form lipoplexes and polyplexes (Lipofectamine 3000 and, respectively, Mirus *Trans*IT). The transfection reagent working solutions were prepared following the manufacturer’s guidelines for the recommended ratio of transfection reagent to DNA. All samples were characterized using a Nano Tracking Assay (NTA) and described in terms of concentration and population size. According to the NTA analysis, Lipofectamine 3000 and Mirus *Trans*IT samples prepared at the recommended concentration for successful transfections have a median size of 220 ± 7 nm (4.5 × 10^8^ ± 3.3 × 10^6^ particles/mL), respectively 197 ± 5 nm (1.0 × 10^8^ ± 3 × 10^7^ particles/mL).

### TRPA1 activity modulation by intracellular Ca^2+^

To determine the effects of our nanocarrier preparations on TRPA1, we adapted the Ca^2+^-imaging setup to more closely reflect physiological conditions, using slow particle perfusion followed by no-flow conditions at physiological temperature. Upon Krebs injection, CHO-mTRPA1 cells displayed TRPA1-dependent Ca^2+^ activity characterized by shallow, undulatory baselines. TRPA1 can be activated by pore-permeating Ca^2+^ following stochastic channel opening [23]. Local Ca^2+^ microdomains amplify channel activity until elevated Ca^2+^ promotes rapid inactivation, after which cytosolic Ca^2+^ is restored through uptake into intracellular stores or extrusion from the cell. As TRPA1 is also temperature- and mechanosensitive, we compared basal activity at room and physiological temperatures under constant flow and no-flow conditions. The conditions that we considered as being most physiologically relevant (i.e. at 35 °C and under no flow) foster the largest basal Ca^2+^ activity in CHO-mTRPA1 cells.

[24][25,26][27][28,29][30][31–33]The interaction between TRPA1 and Ca^2+^ is complex, involving both direct Ca^2+^ binding and Ca^2+^-dependent interactions with calmodulin (CaM) [24]. Although reported EC_50_ values for TRPA1 activation by intracellular Ca^2+^ range from 900 nM to 6 μM [25–27], these concentrations may be reached locally within Ca^2+^ microdomains formed near open channels. Electrophysiological studies show that Ca^2+^ influx leads to peak TRPA1 activation within 30–60 seconds, followed by inactivation within approximately 2 min [28,29], consistent with the 1.30 ± 0.06 min average duration of basal Ca^2+^ transients observed here. CaM further contributes to this self-regulation: Ca^2+^-dependent CaM binding promotes TRPA1 activation at lower Ca^2+^ levels, but desensitization at higher concentrations [30–33]. Thus, local interactions between TRPA1, Ca^2+^ and CaM provide a self-regulating mechanism in which Ca^2+^ influx initially promotes channel activity, but subsequently drives desensitization.

### Cellular uptake of nanocarriers

Immediately after injection, all nanocarrier preparations induced an atypical activation pattern, characterized by one or multiple Ca^2+^ transients of variable amplitude distributed throughout the 12-min measurement window. This contrasts with the smoother, sustained response typically observed with TRP agonists, followed by slower decay during channel desensitization. The atypical pattern may result from stochastic nanocarrier–cell interactions, whereby individual particles or groups of particles induce transient membrane perturbations during internalization.

Thermo Fisher describes uptake of cationic lipid- and polymer-based nanocarriers as driven primarily by endocytosis [34,35]. Previous studies of similarly sized Lipofectamine 2000 lipoplexes (218 nm) found uptake to involve heparan sulfate proteoglycans and the actin cytoskeleton, with only a minor contribution from direct membrane fusion [36]. Comparable mechanisms have been reported for the cationic polymer-based reagent DharmaFECT1 [37]. Lipofectamine 2000 uptake has also been detected within 5 min by live fluorescence imaging [38], while internalized SAINT-2 and DOPE cationic lipoplexes have been observed by electron microscopy within 10 min of application [39] [40]. Thus, the timing of reported nanocarrier uptake is consistent with our Ca^2+^ transients arising during nanocarrier internalization.

### The role of the physiological microenvironment on nanocarrier properties

An important difference between our experimental conditions and physiological nucleic acid delivery is the presence of a biomolecular corona around nanocarriers. In typical *in vitro* transfection protocols, lipid/DNA or polymer/DNA complexes are formed in serum-depleted medium before being transferred to serum-containing medium. Serum proteins can alter nanocarrier uptake pathways: lipoplexes have been reported to undergo clathrin-dependent endocytosis in the absence of serum albumin, whereas albumin favors caveolae-dependent uptake [41]. Similarly, serum can increase membrane fusion by reducing the proportion of particles remaining membrane-bound [42]. These findings indicate that the cellular effects observed in our simplified experimental system may differ in the presence of serum and other components of the physiological microenvironment.

### TRPA1 is partly involved in nanocarrier-induced Ca^2+^ transients

Within one minute of injection, all tested cationic nanocarriers induced Ca^2+^ transients in target cells. Channel blockers reduced but did not fully abolish these responses, indicating that TRPA1 contributes only partially to nanocarrier-induced Ca^2+^ signaling. For Lipofectamine 3000, RR almost completely suppressed responses during the first half of the measurement window, whereas smaller responses persisted later. The different effects of specific and non-specific blockers on the temporal distribution of Ca^2+^ transients may indicate that distinct mechanisms contribute to the early and late responses.

In CHO-mTRPA1 cells, Lipofectamine 3000 and Mirus *Trans*IT produced J- and bell-shaped dose–response curves, respectively. Lipofectamine inhibits TRPA1 at low concentrations, while higher concentrations induce cellular activation, producing the observed J-shaped response. Mirus may similarly exert opposing effects, with activation at lower and inhibition at higher concentrations. Bimodal concentration-dependent effects have also been reported for TRPA1 modulators such as menthol, thymol, cinnamaldehyde, and camphor, which activate the channel at low concentrations but inhibit it at higher concentrations [43,44]. LPS produces a similar pattern and appears to activate TRPA1 by altering plasma membrane organization rather than through classical ligand–channel binding [11]. Thus, the relative balance between concentration-dependent activation and inhibition can produce J-, U-, or bell-shaped overall responses.

### TRPA1-independent generation of Ca^2+^ transients

As none of the TRPA1 blockers fully abolished nanocarrier-induced Ca^2+^ responses, we confirmed TRPA1-independent signaling in CHO-WT cells. Cationic lipo- and polyplexes induced similar, albeit weaker, Ca^2+^ transients in wild-type cells, which were partially inhibited by the non-specific Ca^2+^ channel blocker RR. Because RR cannot cross the plasma membrane due to its large size and positive charge, this indicates that at least part of the response results from extracellular Ca^2+^ influx, either through other Ca^2+^-permeable ion channels or through de novo pore formation caused by membrane destabilization [45].

### The role of polyamines in nanocarrier-target cell interactions

Cationic lipids and polymers contain amine groups that confer positive charge at physiological pH. These chemical groups may contribute to nanocarrier–cell interactions and potentially influence ion channel activity. Natural polyamines such as spermine, spermidine, and putrescine can modulate multiple TRP channels, with effects depending on concentration and channel type. Polyamines can activate and sensitize TRPV1 at millimolar concentrations, but inhibit several TRPV and TRPC channels at lower concentrations [46–49]. They also inhibit members of the TRPM family, with potency depending on the net positive charge of the molecule [50,51]. This inhibition appears to involve pore permeation and interactions with negatively charged residues within the channel pore [2]. Polyamines can also bind the intracellular lipid PIP_2_, potentially reducing activation of PIP_2_-dependent channels such as TRPA1 [53]. In addition, tertiary amine-containing local anesthetics such as lidocaine produce concentration-dependent bimodal effects on TRPA1, activating the channel at low concentrations and inhibiting it at higher concentrations through a membrane-permeable mechanism [54].

Although there is no direct evidence that cationic lipids or polymers modulate TRPA1, these findings demonstrate that positively charged amine-containing compounds can exert complex, concentration-dependent effects on ion channels. It is therefore plausible that the amine-containing components of nanocarriers contribute to both activation and inhibition of TRPA1, while also affecting other ion channels. This could help explain the partial contribution of TRPA1 to the Ca^2+^ responses observed in our experiments.

### Nanocarriers induce Ca^2+^ transients in sensory neurons

Lastly, we investigated lipid-based nanocarriers in a native TRPA1-expressing system using primary sensory DRG neurons. Both cationic lipoplexes induced transient Ca^2+^ responses similar to those observed in CHO-mTRPA1 cells, although responses were substantially smaller. Ca^2+^ mobilization from intracellular stores was also minimal, potentially reflecting differences in Ca^2+^ handling and the relative contribution of IP_3_R- and RyR-mediated release pathways between CHO cells and sensory neurons [30,55,56]. We further assessed TRPA1 involvement in Lipofectamine 3000-induced responses. HC-030031 and RR significantly reduced Lipo 10 responses across all parameters, whereas Lipo 5 responses were unaffected by HC-030031 but reduced by RR. These findings indicate partial, concentration-dependent TRPA1 involvement alongside additional TRPA1-independent mechanisms.

DRG neurons are heterogeneous and express numerous ion channels and signaling receptors, providing several potential targets for lipoplex-induced Ca^2+^ signaling. In addition to the polyamine-sensitive channels discussed above, TRPC channels can be activated downstream of GPCR/PLC signaling or by increased intracellular Ca^2+^ [1,57]. Cationic lipids have also been reported to interact with membrane proteins including integrins, TLR4, GPCRs, and MD-2. Some cationic or ionizable lipids share structural features with the lipid A component of LPS, including amphipathic geometry, multiple hydrophobic chains, and the ability to interact with anionic structures and alter membrane organization. Such similarities could enable related signaling effects despite their different net charges [58]. Importantly, both the cationic headgroup and hydrophobic tails appear necessary for cationic lipids to induce PLC-dependent Ca^2+^ release, suggesting that the overall molecular architecture of the nanocarrier, rather than charge alone, contributes to Ca^2+^ homeostasis [59].

### Effects of nanocarriers on cell signaling and cellular function

Although this study focuses on cationic liposome-based transfection reagents for *in vitro* application, cationic lipoplexes are also widely used for *in vivo* nucleic acid delivery. In both settings, the permanent positive charge of cationic lipoplexes and polyplexes has been associated with increased cytotoxicity [60], which can correlate with transfection efficiency. For Lipofectamine 3000, transfection efficiency is substantially higher when P3000 is included, but this is accompanied by reduced cell viability [61]. Consistent with this, our single-component experiments showed that Lipofectamine complexes had little effect on Ca^2+^ homeostasis, whereas Lipofectamine:P3000 complexes produced clear alterations, including inhibition of basal Ca^2+^ activity at low concentrations in CHO-mTRPA1 cells and induction of Ca^2+^ activity at higher concentrations in CHO-WT cells. These findings raise the possibility that nanocarrier-induced Ca^2+^ signaling contributes to the cytotoxic effects of cationic lipo- and polyplexes, which warrants further investigation.

Ca^2+^ is a key second messenger involved in numerous cellular signaling processes, with the spatiotemporal properties of Ca^2+^ signals determining diverse physiological outcomes. Ca^2+^ spikes can trigger exocytosis, signaling factor release, cell differentiation, proliferation, transcription factor activation, and apoptosis, and can regulate gene expression more effectively than sustained signals of similar average amplitude [24,62]. Cytosolic Ca^2+^ also regulates the expression of plasma membrane ion channels and pumps, including TRPC channels [24,63]. Conversely, large or sustained elevations in intracellular Ca^2+^ can have deleterious effects, including activation of Ca^2+^-dependent proteases, reactive oxygen species (ROS) production, organelle remodeling, and mitochondrial permeability transition. Mitochondria are particularly susceptible because of their proximity to major Ca^2+^ sources such as the ER and plasma membrane [56,63].

Cationic lipids have been reported to induce pro-apoptotic and pro-inflammatory responses, including ROS production, chemokine and cytokine secretion, and expression of co-stimulatory and maturation markers. Lipofectamine-induced Ca^2+^ release from intracellular stores may also contribute to ROS production. Through their tertiary and quaternary amine groups, cationic lipids can modulate kinases and proteases such as PKC, MAPK, and caspases and activate NF-κB, apoptotic, and inflammasome pathways. These properties can be beneficial for the immunostimulatory activity and adjuvant function of cationic lipoplexes *in vivo*. Cationic lipids have also been reported to interact with CD14, promoting β-integrin clustering and downstream PLC activation, which may contribute to Ca^2+^ mobilization from intracellular stores [58,60,64]. Given these reported effects on cellular signaling and function, nanocarriers capable of acutely perturbing cytosolic Ca^2+^ homeostasis should therefore be considered carefully, particularly when intended for therapeutic applications.

To conclude, here we show that nanocarriers based on cationic lipids and cationic polymers are able to induce irregular Ca^2+^ transients in target cells, both dependently and independently of TRPA1. However, the effect in CHO-mTRPA1 cells are significantly more pronounced than in CHO-WT cells. We envisage that nanocarriers induce a multifactorial effect in CHO-mTRPA1 cells. Firstly, they induce Ca^2+^ mobilization from internal stores, in particular the endoplasmic reticulum. Secondly, they trigger Ca^2+^ influx across the plasma membrane. Part of the Ca^2+^ influx is reduced in the presence of TRPA1 channel blockers, indicating a partial contribution of TRPA1 to the observed Ca^2+^ transients. The TRPA1-mediated Ca^2+^ influx is likely downstream of Ca^2+^ mobilization from the ER and acts in amplifying the Ca^2+^ release from internal stores. Other direct effects, such as TRPA1 activation induced by plasma membrane fluidization or direct activation by trace free lipids and/or polymers in the preparations could also contribute to the TRPA1-dependent Ca^2+^ responses. The molecular mechanisms underlying the remaining TRPA1-independent Ca^2+^ influx are not yet identified, and more work is required in order to better understand the multifactorial effects of lipo- and polyplexes. The nanoparticles tested here may engage other players at the plasma membrane level besides TRPA1, such as other ion channels and G-protein coupled receptors, or could induce Ca^2+^-permeable pores in the plasma membrane. Nonetheless, here we have identified a new class of TRPA1 activity modulators, lipid- and polymer-based nanoparticles, that induce a non-canonical pattern of activation. These results serve as a starting point in better understanding the interplay between TRP channels and various lipid particles involved in physiological and pathophysiological processes in the body.

## Acknowledgements

We thank Melissa Benoit for the technical support and the members of the LICR for helpful discussions. The CHO-mTRPA1 cell line was kindly provided by Ardem Patapoutian (The Scripps Research Institute, USA).

## Author Contributions

All authors contributed to the study conception and design. Material preparation, data collection and analysis were performed by Alina Milici. Andrei Segal designed and built the thermoregulated system for the Ca^2+^ imaging experiments. The first draft of the manuscript was written by Alina Milici and Karel Talavera and all authors commented on it. All authors read and approved the final manuscript. The Python script for the detection of Ca^2+^ transients was developed in collaboration with Tamas Barath (KBC, Department of Advanced Data Analytics and Modelling, Belgium).

## Data Availability

All data generated or analyzed during this study are included in the supporting files and online in the Research Data Repository of KU Leuven. The Python script used to identify the Ca^2+^ transients is available on GitHub [21].

## Ethics approval

The mouse tissue harvest was done in accordance with the European Community Council guidelines and were approved was approved by the Ethical Committee for Animal Experimentation (ECD) -KU Leuven/UZ Leuven.

